# Integration of electric, calcium, reactive oxygen species and hydraulic signals during rapid systemic signaling in plants

**DOI:** 10.1101/2021.02.12.430927

**Authors:** Yosef Fichman, Ron Mittler

**Author notes:** Corresponding author: Ron Mittler **Email:**.

## Abstract

The sensing of abiotic stress, mechanical injury, or pathogen attack by a single plant tissue results in the activation of systemic signals that travel from the affected tissue to the entire plant, alerting it of an impending stress or pathogen attack. This process is essential for plant survival during stress and is termed systemic signaling. Among the different signals triggered during this process are calcium, electric, reactive oxygen species (ROS) and hydraulic signals. These are thought to propagate at rapid rates through the plant vascular bundles and to regulate many of the systemic processes essential for plant survival. Although the different signals activated during systemic signaling are thought to be interlinked, their coordination and hierarchy remain to be determined. Here, using a combination of advanced whole-plant imaging and hydraulic pressure measurements, we studied the activation of all four systemic signals in wild type and different *Arabidopsis thaliana* mutants subjected to a local high light (HL) stress or wounding. Our findings reveal that in response to wounding systemic changes in membrane potential, calcium, ROS, and hydraulic pressure are coordinated by glutamate receptor-like (GLR) proteins 3.3 and 3.6, while in response to HL the respiratory burst oxidase homolog D-driven systemic ROS signal could be separated from systemic changes in membrane potential and calcium levels. We further determine that plasmodesmata functions are required for systemic changes in membrane potential, calcium, and ROS during systemic signaling. Our findings shed new light on the different mechanisms that integrate different systemic signals in plants during stress.

**Significance statement:** The ability of plants to transmit a signal from a stressed or wounded tissue to the entire plant, termed systemic signaling, is key to plant survival during conditions of environmental stress. At least four different systemic signals are thought to be involved in this process: electric, calcium, reactive oxygen and hydraulic. However, how are they coordinated and whether they can be stress-specific is mostly unknown. Here we report that different types of stimuli can induce different types of systemic signals that may or may not be linked with each other. We further reveal that hydraulic waves can be actively regulated in plants in response to wounding, and that proteins that regulate plasmodesmata pores play a key role in systemic signaling.

## Introduction

Being sessile organisms, plants evolved multiple defense and acclimation mechanisms that enable them to rapidly respond to different stimuli and/or stress conditions in their environment (1–4). Changes in light intensity, temperature, humidity, and/or herbivore, or pathogen attack, for example, activate within seconds different signal transduction pathways that regulate different molecular, metabolic, and physiological responses, critical for plant survival during stress (4, 5). The sensing of stress, pathogen attack and/or mechanical injury not only induces defense and acclimation responses at the affected plant tissue(s), but also triggers a rapid systemic signal transduction process that alerts all other parts of the plant to the impending change in the environment and/or a pathogen/herbivore attack (6–11). This process was shown to occur in multiple plant species in response to many different stimuli or stress conditions and is termed systemic signaling. Upon perception of the systemic signal (that was generated at the affected parts of the plant), different molecular, metabolic and physiological responses are activated in systemic tissues, resulting in a heightened whole-plant state of systemic acquired acclimation (SAA; 12) to abiotic stress, systemic acquired resistance (SAR; 13) to pathogen attack, and/or systemic wound response (SWR; 14) to mechanical injury or herbivore attack.

Because plants lack a nervous system that connects the sensing (*i.e.,* local) tissue with all other plant tissues that did not yet experience the stress/pathogen/stimuli (*i.e.,* systemic tissues), systemic signals in plants must travel from cell-to-cell over long distances, sometimes spanning the entire length of the plant (6–11). Among the different signaling processes thought to mediate such rapid long distance cell-to-cell signal transduction mechanisms in plants are changes in membrane potentials (*i.e.,* electric waves; 7, 15–19), steady-state levels of calcium (*i.e.,* calcium wave; 8, 18, 20–24), steady-state levels of reactive oxygen species (ROS; *i.e.,* ROS wave; 4, 6, 9, 25–30), hydraulic pressure (*i.e.,* hydraulic wave; 31–33), as well as rapid changes in the levels of different plant hormones such as jasmonic acid (JA), abscisic acid (ABA) and auxin (34–38), small peptides (39), redox levels (40), and/or different metabolites/metabolic signatures (41). Recent studies demonstrated that many of these signals propagate from cell-to-cell through the vascular bundles of plants using tissues such as xylem parenchyma and phloem cells to mediate systemic electric, calcium and ROS signals, and xylem cells to mediate hydraulic pressure signals (16–18, 21, 28, 31, 42–44). In addition, the calcium channels glutamate receptor-like (GLR) 3.3 and 3.6 were found to play a key role in regulating systemic electric and calcium signals during wounding (16–18, 21), and the respiratory burst oxidase homolog D and F (RBOHD and RBOHF) proteins were found to play a key role in the regulation of systemic ROS signals during high light (HL) stress (25–29). Systemic ROS signals were further shown to be regulated by cyclic nucleotide-gated calcium channel 2 (CGNC2), mechanosensitive small conductance–like (MSL) channels 2 and 3, and plasmodesmata (PD)-localized proteins (PDLP) 1 and 5, during systemic responses to HL stress (44).

Although many of the rapid systemic signaling mechanisms described above were characterized, and the proteins underling some of them identified, it is unknown at present how they interact with each other, and how they respond to different stimuli (6–8, 10, 11). Moreover, studies identifying and characterizing each of the different systemic signals described above were conducted in different laboratories around the world using plants grown under different growth conditions, subjected to different types and degrees of stress treatments, as well as measured using different methods and equipment. To study the integration of different systemic signals, their specificity, and their hierarchical-/co-regulation, under similar experimental conditions, we measured systemic changes in membrane potential, calcium and ROS levels, as well as hydraulic pressure, in 4-5 week-old *Arabidopsis thaliana* wild type, *rbohD, glr3.3;glr3.6* and *pdlp5* plants grown under the same growth conditions and subjected to the same local treatments of HL stress or wounding. Our findings reveal that in response to wounding, systemic changes in membrane potential, calcium, ROS and hydraulic pressure are coordinated by GLR3.3;GLR3.6, while in response to HL stress, RBOHD-mediated systemic ROS signals could be separated from GLR3.3;GLR3.6-mediated changes in membrane potential and calcium levels. We further identify a novel dependency of systemic hydraulic pressure signals on GLR3.3;GLR3.6 function during wounding (but not HL stress), and determine that PD functions are required for systemic changes in membrane potential, calcium, and ROS levels, during systemic responses to HL stress or wounding. Our findings reveal that different systemic signaling mechanisms and pathways could be activated by different stresses, that many of these systemic signaling pathways are interlinked, and that many of them require the function of PD-associated proteins.

## Results

### Whole-plant changes in ROS levels in wild type, *rbohD, glr3.3;glr3.6* and *pdlp5* plants subjected to a local treatment of HL stress or wounding

To follow changes in local and systemic ROS levels in response to a local treatment of HL stress or wounding in wild type plants and the different mutants, we used the whole-plant live ROS imaging method we developed and used to study systemic ROS signals (26). Using this method, we were able to determine that the systemic ROS signal propagates through the vascular bundles in response to HL or wounding, and through the vascular bundles and/or mesophyll cells in response to wounding or heat stress (28, 42), that in response to HL the systemic ROS signal requires the function of RBOHD, PDLP5, CNGC2, plasma membrane intrinsic protein (PIP) 2;1, MSL2 and other proteins (44), and that during a combination of two different stresses applied to two different leaves, the systemic ROS signal is involved in integrating the two different systemic signals generated by the two different stresses (27). In agreement with our previous studies (44), in response to a local application of HL stress, the systemic ROS signal was blocked in the *rbohD* and *pdlp5* mutants, but only suppressed in the *glr3.3;glr3.6* double mutant (Fig. 1, *SI Appendix*, Fig. S1). In contrast, in response to wounding, the systemic ROS signal was blocked in all three mutants tested *(rbohD, glr3.3;glr3.6, and pdlp5;* Fig. 1). These findings suggest that the regulation of the systemic ROS signal in response to HL is different than that in response to wounding. In response to wounding the systemic ROS signal is dependent on the GLR3.3;GLR3.6 proteins, while in response to HL it is not (Fig. 1; 44).

**Fig. 1.**
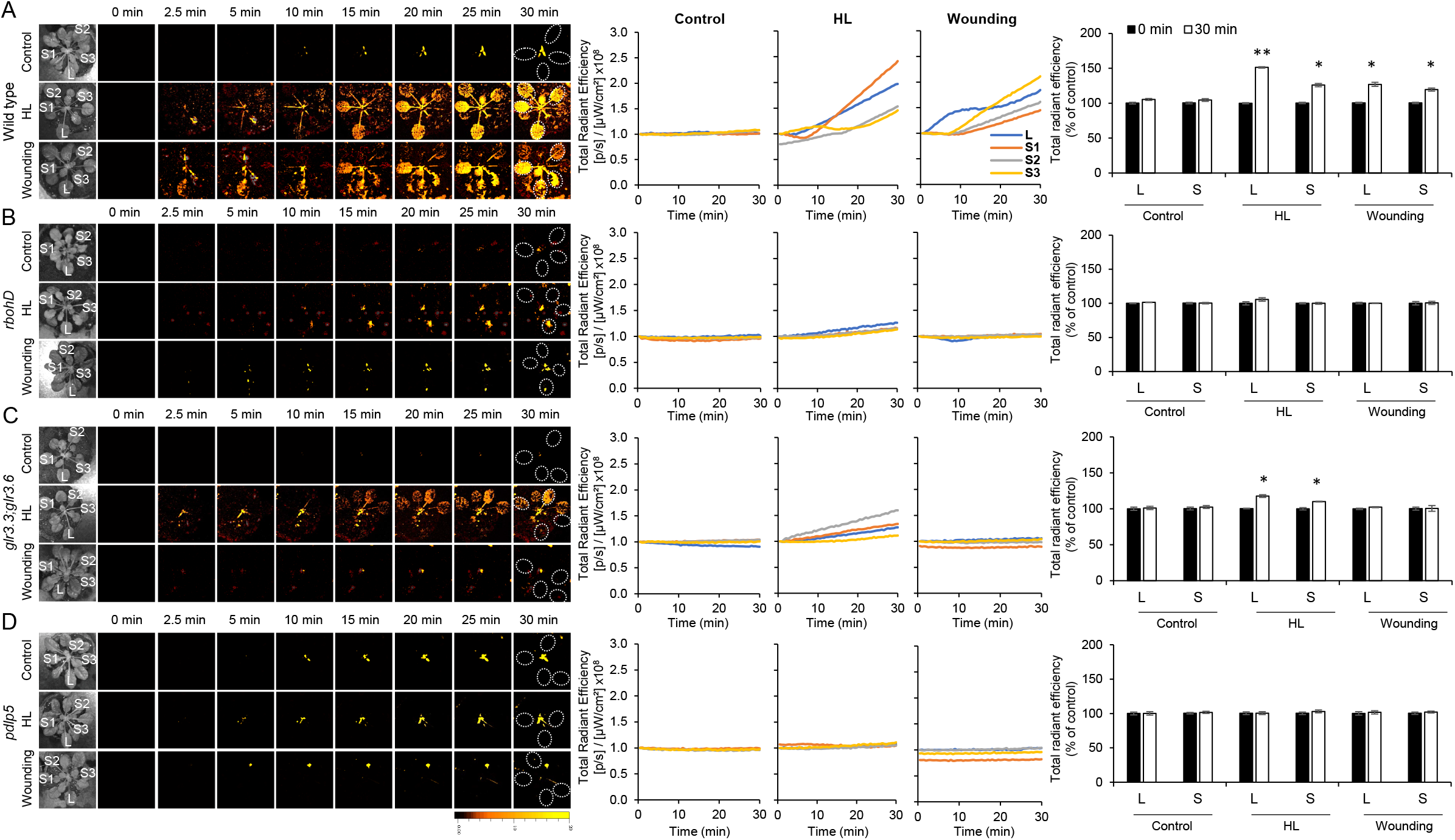
Whole-plant changes in ROS levels in wild type, *rbohD*, *glr3.3;glr3.6* and *pdlp5* plants subjected to a local wounding or high light stress treatment. (*A*) Time-lapse imaging of whole-plant ROS accumulation in untreated (control) wild type *Arabidopsis thaliana* plants, and wild type plants subjected to a 2 min local (L) high light (HL) stress or wounding treatment (applied to leaf L only) is shown on left; continuous measurements of ROS levels in the local (L) and systemic (S1-S3) leaves over the entire course of the experiment are shown in the middle (ROIs used for calculating them are indicated with white dotted ovals on the images to the left); and statistical analysis of ROS accumulation in local and systemic leaves at 0 and 30 min of untreated (control), HL stress, or wounding treatments is shown on right. (*B*) Same as (*A*) but for *rbohD*. (*C*) Same as (*A*) but for *glr3.3;glr3.6*. (*D*) Same as (*A*) but for *pdlp5*. All experiments were repeated at least 3 times with 10 plants per biological repeat. Student t-test, SE, N=30, *p < 0.05, **p < 0.01. Scale bar indicates 1 cm. *Abbreviations used*: glr, glutamate receptor-like; HL, high light; L, local; pdlp5, plasmodesmata localized protein 5; rbohD, respiratory burst oxidase homolog D; ROI, region of interest; S, systemic.

### Whole-plant changes in calcium levels in wild type, *rbohD, glr3.3;glr3.6* and *pdlp5* plants subjected to a local treatment of HL stress or wounding

To follow changes in local and systemic calcium levels in response to a local treatment of HL stress or wounding in wild type plants and the different mutants, we used the same application method used in our whole-plant live ROS imaging method (Fig. 1, *SI Appendix*, Fig. S1; 26–28), but instead of applying dichlorofluorescein (DCF) as H_2_DCFDA by fumigation, we applied the Fluo-4-AM calcium-sensitive dye. This dye was previously used to measure changes in cytosolic calcium levels in plant and animal cells (45–48), and in control experiments it was responsive to a combined treatment of calcium and a calcium ionophore (*SI Appendix*, Fig. S2). Changes in calcium levels, measured as increased fluorescence of the Fluo-4-AM dye, occurred in local and systemic leaves of wild type plants subjected to a local treatment of HL stress or wounding (Fig. 2). These measurements indicated that the local application of HL stress or wounding resulted in a systemic increase in calcium levels, similar to the induction of the cytosolic calcium wave, previously reported in response to a local treatment of wounding or salinity stress in Arabidopsis (8, 21, 23, 24). In contrast, similar changes were not observed in *rbohD, glr3.3;glr3.6* and *pdlp5* plants in response to a local application of HL stress or wounding (Fig. 2). These findings suggest that in response to a local HL stress or wounding, systemic changes in calcium levels are dependent on the RBOHD, GLR3.3;GLR3.6 and PDLP5 proteins. Previous studies revealed that in response to wounding, the calcium wave was dependent on GLR3.3;GLR3.6 (21), and in response to salinity, on two-pore channel (TPC) 1 and RBOHD (23, 24), supporting the validity of our results with the Fluo-4-AM dye (Fig. 2, *SI Appendix*, Fig. S2). Our current analysis therefore reveals that in response to wounding, systemic calcium and ROS signals are linked and require RBOHD, GLR3.3;GLR3.6 and PDLP5, while in response to HL stress, the systemic ROS signal could be separated from the systemic calcium signal and occur in the *glr3.3;glr3.6* mutant that does not show a systemic calcium signal with the Fluo-4-AM dye (Figs. 1, 2; 44).

**Fig. 2.**
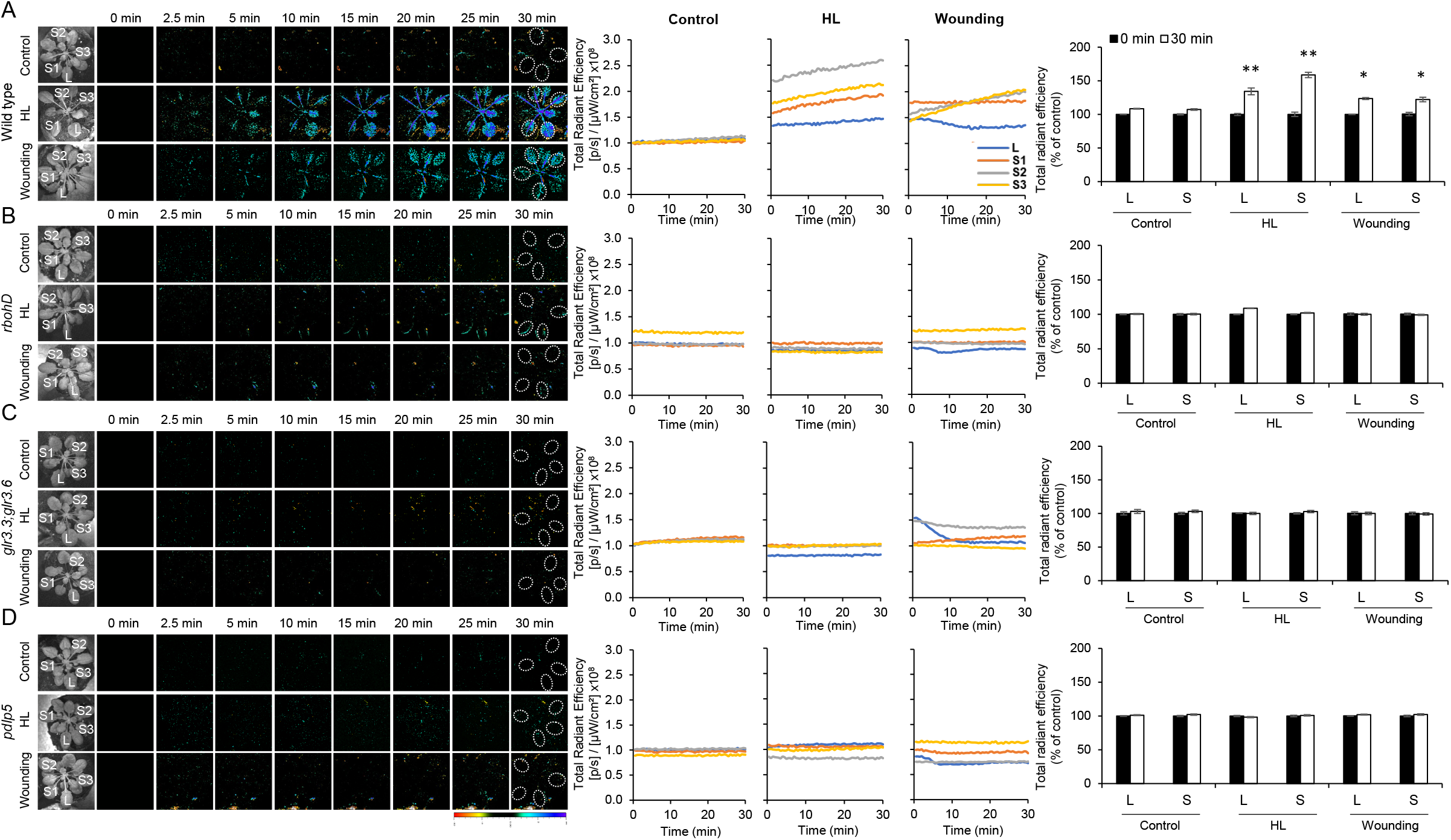
Whole-plant changes in calcium levels in wild type, *rbohD, glr3.3;glr3.6* and *pdlp5* plants subjected to a local wounding or high light stress treatment. (*A*) Time-lapse imaging of whole-plant changes in calcium levels in untreated (control) wild type *Arabidopsis thaliana* plants, and wild type plants subjected to a 2 min local (L) high light (HL) stress or wounding treatment (applied to leaf L only) is shown on left; continuous measurements of changes in calcium levels in the local (L) and systemic (S1-S3) leaves over the entire course of the experiment are shown in the middle (ROIs used for calculating them are indicated with white dotted ovals on the images to the left); and statistical analysis of changes in calcium levels in local and systemic leaves at 0 and 30 min of untreated (control), HL stress, or wounding treatments is shown on right. (*B*) Same as (*A*) but for *rbohD*. (*C*) Same as (*A*) but for *glr3.3;glr3.6*. (*D*) Same as (*A*) but for *pdlp5*. All experiments were repeated at least 3 times with 10 plants per biological repeat. Student t-test, SE, N=30, *p < 0.05, **p < 0.01. Scale bar indicates 1 cm. *Abbreviations used:* glr, glutamate receptor-like; HL, high light; L, local; pdlp5, plasmodesmata localized protein 5; rbohD, respiratory burst oxidase homolog D; ROI, region of interest; S, systemic.

### Whole-plant changes in membrane potential in wild type, *rbohD, glr3.3;glr3.6* and *pdlp5* plants subjected to a local treatment of HL stress or wounding

To measure local and systemic changes in membrane potential in response to a local treatment of HL stress or wounding in wild type plants and the different mutants, we used the same application method used in Figs. 1, 2, and *SI Appendix*, Figs. S1, S2, but instead of applying H_2_DCFDA or Fluo-4-AM by fumigation, we applied the membrane potential sensitive dye DiBAC4(3) [Bis-(1,3-Dibutylbarbituric Acid)Trimethine Oxonol]. This dye was previously used to measure changes in membrane potential in different plant cells and tissues, as well as action and variation potentials (AP and VP, respectively), two electric wave forms that lead to SWR and/or SAA, in vascular bundles and phloem cells (49–52). In control experiments DiBAC4(3) was responsive to a treatment with sodium cholate (49) that altered membrane potentials (*SI Appendix*, Fig. S3). Changes in membrane potential, measured as increased fluorescence of the DiBAC4(3) dye, occurred in local and systemic leaves of wild type plants subjected to a local treatment of HL or wounding (Fig. 3). These measurements indicate that the local application of HL stress or wounding resulted in the triggering of systemic membrane potential changes that spread through the entire plant.

**Fig. 3.**
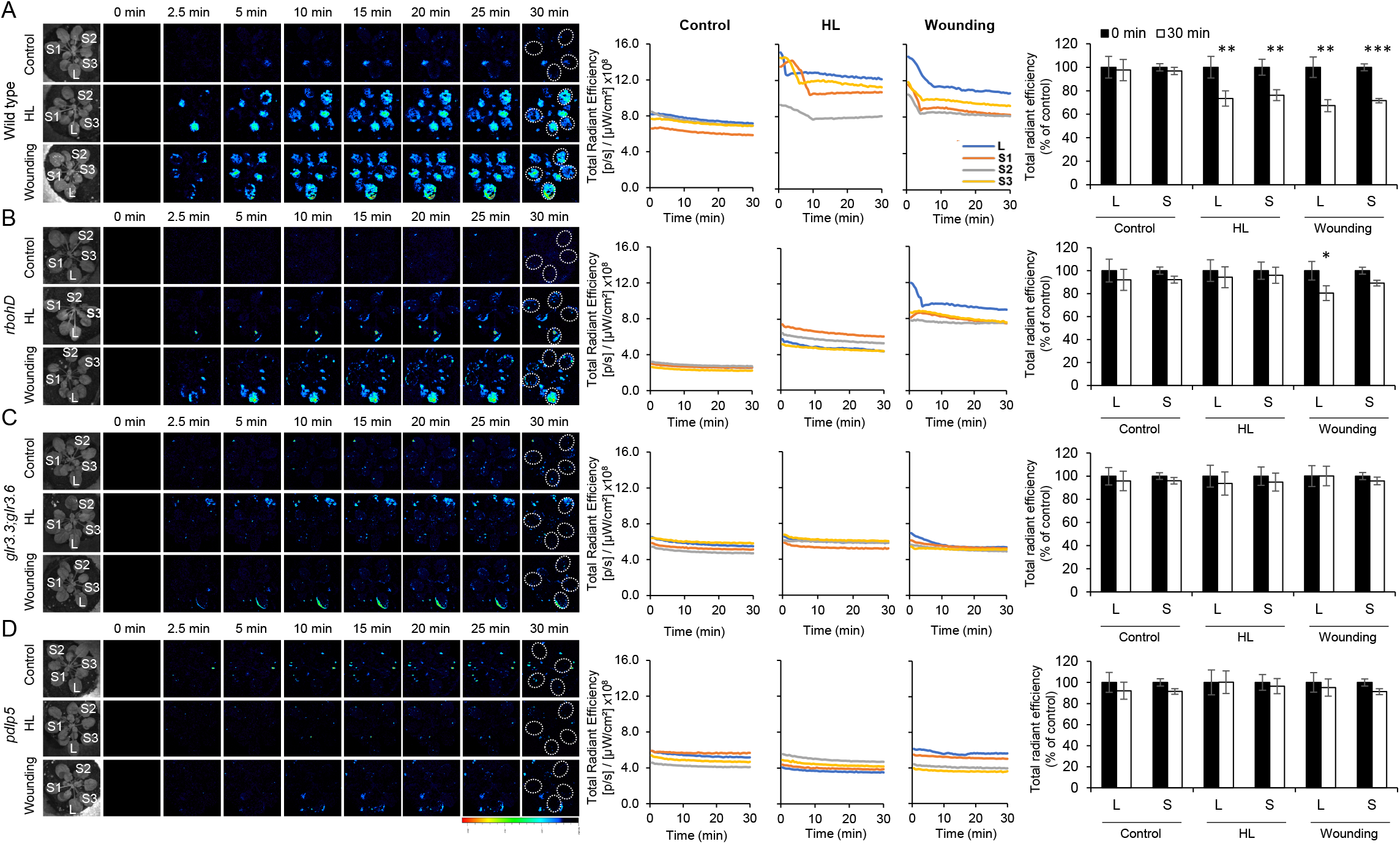
Whole-plant changes in membrane potential in wild type, *rbohD, glr3.3; glr3.6* and *pdlp5* plants subjected to a local wounding or high light stress treatment. (*A*) Time-lapse imaging of whole-plant changes in membrane potential in untreated (control) wild type *Arabidopsis thaliana* plants, and wild type plants subjected to a 2 min local (L) high light (HL) stress or wounding treatment (applied to leaf L only) is shown on left; continuous measurements of changes in membrane potential in the local (L) and systemic (S1-S3) leaves over the entire course of the experiment are shown in the middle (ROIs used for calculating them are indicated with white dotted ovals on the images to the left); and statistical analysis of changes in membrane potential in local and systemic leaves at 0 and 30 min of untreated (control), HL stress, or wounding treatments is shown on right. (*B*) Same as (*A*) but for *rbohD*. (*C*) Same as (*A*) but for *glr3.3;glr3.6*. (*D*) Same as (*A*) but for *pdlp5*. All experiments were repeated at least 3 times with 10 plants per biological repeat. Student t-test, SE, N=30, *p < 0.05, **p < 0.01, ***p < 0.001. Scale bar indicates 1 cm. *Abbreviations used*: glr, glutamate receptor-like; HL, high light; L, local; pdlp5, plasmodesmata localized protein 5; rbohD, respiratory burst oxidase homolog D; ROI, region of interest; S, systemic.

APs and VPs were previously reported in response to a local treatment of wounding or HL stress in Arabidopsis, supporting the results presented here (7, 15–19). In contrast to the responses measured in wild type plants, systemic changes in membrane potential were not observed in the *rbohD, glr3.3;glr3.6* or *pdlp5* mutants in response to a local application of HL or wounding (Fig. 3). These findings suggest that in response to a local HL stress or wounding, systemic changes in membrane potential are dependent on the RBOHD, GLR3.3;GLR3.6 and PDLP5 proteins. While local membrane potential responses to wounding were suppressed in the *glr3.3;glr3.6* or *pdlp5* mutants, they were not suppressed in the *rbohD* mutant. Previous studies revealed that in response to wounding systemic APs and VPs were dependent on GLR3.3;GLR3.6 (16–18) and in response to HL stress on RBOHD (53), further supporting the validity of our results with the DiBAC4(3) dye (Fig. 3 *SI Appendix*, Fig. S3). Our current analysis therefore reveals that in response to wounding, systemic calcium, membrane potential and ROS levels are linked and require RBOHD, GLR3.3;GLR3.6 and PDLP5, while in response to HL stress the systemic ROS signal can be separated from systemic calcium and membrane potential responses (Figs. 1-3, *SI Appendix*, Figs. S1-S3).

### Systemic changes in hydraulic pressure in wild type, *rbohD, glr3.3;glr3.6* and *pdlp5* plants subjected to a local treatment of HL stress or wounding

To measure systemic changes in hydraulic pressure in wild type plants and the different mutants in response to a local application of HL stress or wounding, we used hydraulic pressure probes developed to monitor leaf water potential and hydraulic pressure in real time in live plants grown in soil (32, 54). In response to a local treatment of HL stress, wild type, as well as the *rbohD, glr3.3;glr3.6* and *pdlp5* mutants displayed a systemic hydraulic pressure signal that initiated almost immediately upon HL stress application and peaked at about 15 min post stress initiation (Fig. 4A, *SI Appendix*, Fig. S4). In contrast, in response to a local wounding treatment, wild type plants displayed a systemic hydraulic pressure signal that initiated immediately upon wounding but did not peak during the first 30 min post wounding (Fig. 4B, *SI Appendix*, Fig. S4). The *rbohD* mutant displayed a systemic hydraulic pressure signal response that was similar to its response to HL, albeit weaker, and the *pdlp5* mutant displayed a systemic hydraulic pressure signal response that was similar to its response to HL, albeit stronger (Fig. 4B). In contrast, the wound-induced systemic hydraulic pressure signal response of the *glr3.3;glr3.6* double mutant was abolished (Fig. 4B). These results suggest that in response to a local HL stress treatment, the systemic hydraulic pressure signal can occur in the absence of the RBOHD, GLR3.3;GLR3.6 or PDLP5 proteins. In contrast, in response to a local wound treatment, the systemic hydraulic pressure signal is dependent on the presence of the GLR3.3;GLR3.6 proteins, and could be amplified in the absence of the PDLP5 protein.

**Fig. 4.**
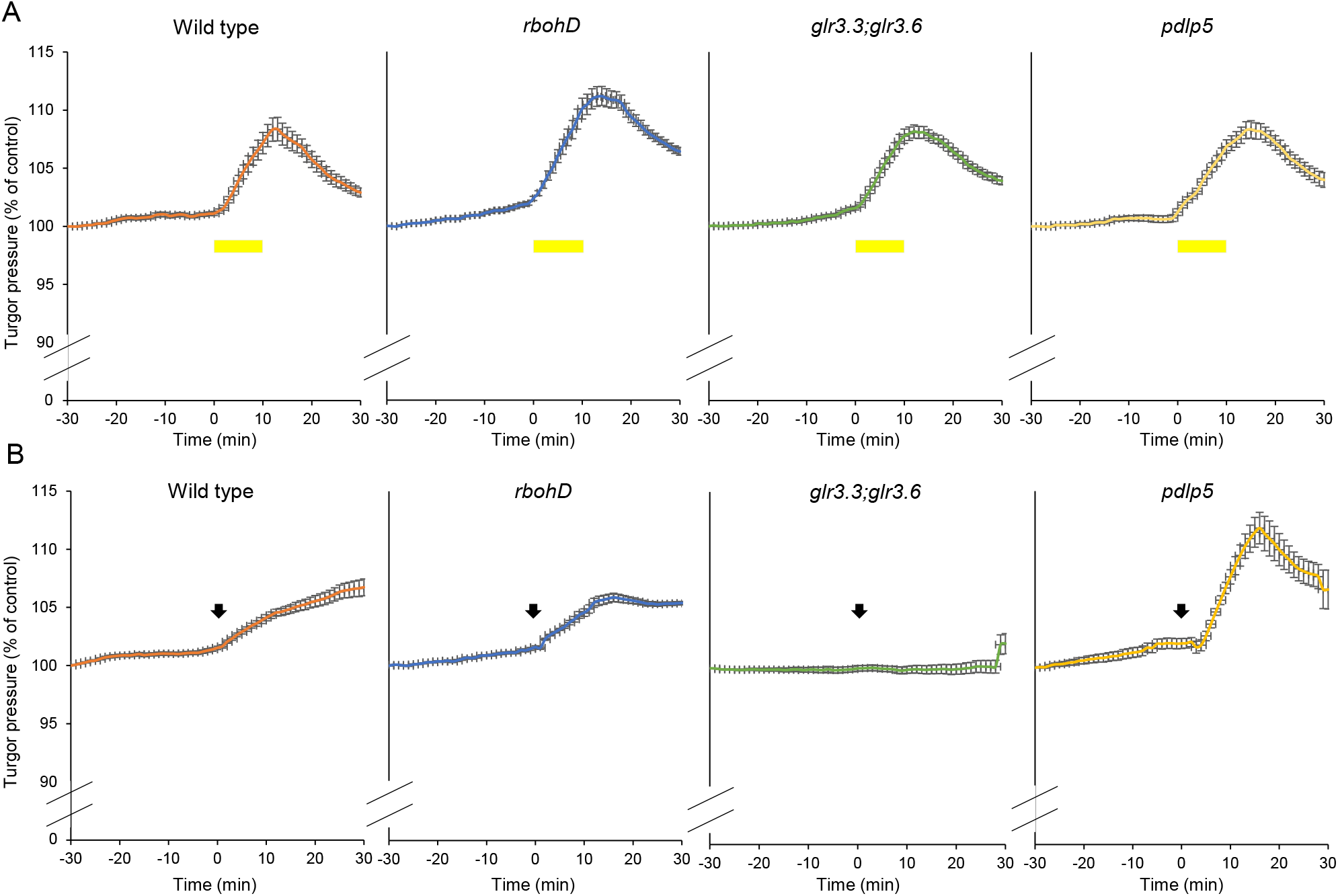
Systemic changes in hydraulic pressure in wild type, *rbohD, glr3.3;glr3.6* and *pdlp5* plants subjected to a local wounding or high light stress treatment. (*A*) Continues systemic leaf turgor pressure measurements of wild type, *rbohD, glr3.3;glr3.6* and *pdlp5* plants, from 30 min prior (−30 min) to 30 min post (30 min) high light (HL) stress treatment (applied from 0-10 min) to a single local leaf. (*B*) Same as (*A*) but for wounding of a local single leaf applied at time 0 min. Hydraulic pressure is represented as percent of the initial measured turgor pressure at −30 min. The period of HL stress application is indicated with a yellow rectangle in (*A*), and the instant of plant wounding is indicated with an arrow in (*B*). All experiments were repeated at least 5 times. SE. *Abbreviations used:* glr, glutamate receptor-like; HL, high light; pdlp5, plasmodesmata localized protein 5; rbohD, respiratory burst oxidase homolog D.

### Local and systemic changes in SAA- and SWR-transcript expression in wild type, *rbohD*, *glr3.3;glr3.6* and *pdlp5* plants subjected to a local HL stress or wounding

To determine whether the systemic signals observed in the different mutants in response to HL stress or wounding (Figs. 1-4) are associated with accumulation of different SAA- and SWR-associated transcripts, we used quantitative real-time PCR (qPCR) to study the expression of *MYB30* and *Zat10*, two HL-induced SAA transcripts (29, 44, 55), and *JAZ5* and *JAZ7*, two wound-induced SWR transcripts (21, 53), in local and systemic leaves of wild type and the different mutants subjected to a local HL or wounding treatment. While expression of *MYB30* and *Zat10* was high in systemic leaves of wild type plants 30 min post HL stress application to a local leaf, the systemic expression of these transcripts was severely suppressed in the different mutants (Fig. 5A). Only in the *glr3.3;glr3.6* double mutant that displayed a suppressed systemic ROS signal response (Fig. 1), some recovery was observed in the systemic expression of *Zat10* (Fig. 5A). This finding was in agreement with our previous findings that in response to a local HL stress treatment, expression of *Zat10* and *MYB30* was only suppressed, but not abolished, at 2 and 8 min in the *glr3.3;glr3.6* double mutant (44). In response to a local treatment of wounding, the systemic expression of *JAZ5* and *JAZ7* was high in wild type plants (Fig. 5B). In contrast, systemic expression of the *JAZ5* and *JAZ7* transcripts was severely suppressed in the different mutants (Fig. 5B). Only in the *rbohD* mutant that displayed suppressed systemic ROS, calcium and electric responses during wounding (Figs. 1-3), some recovery was observed in the systemic expression of *JAZ5* (Fig. 5B). The results presented in Fig. 5 demonstrate that disrupting some of the systemic signals detected in Figs. 1-4 impaired the systemic accumulation of certain SAA- or SWR-transcripts in response to a local HL stress or wounding treatments.

**Fig. 5.**
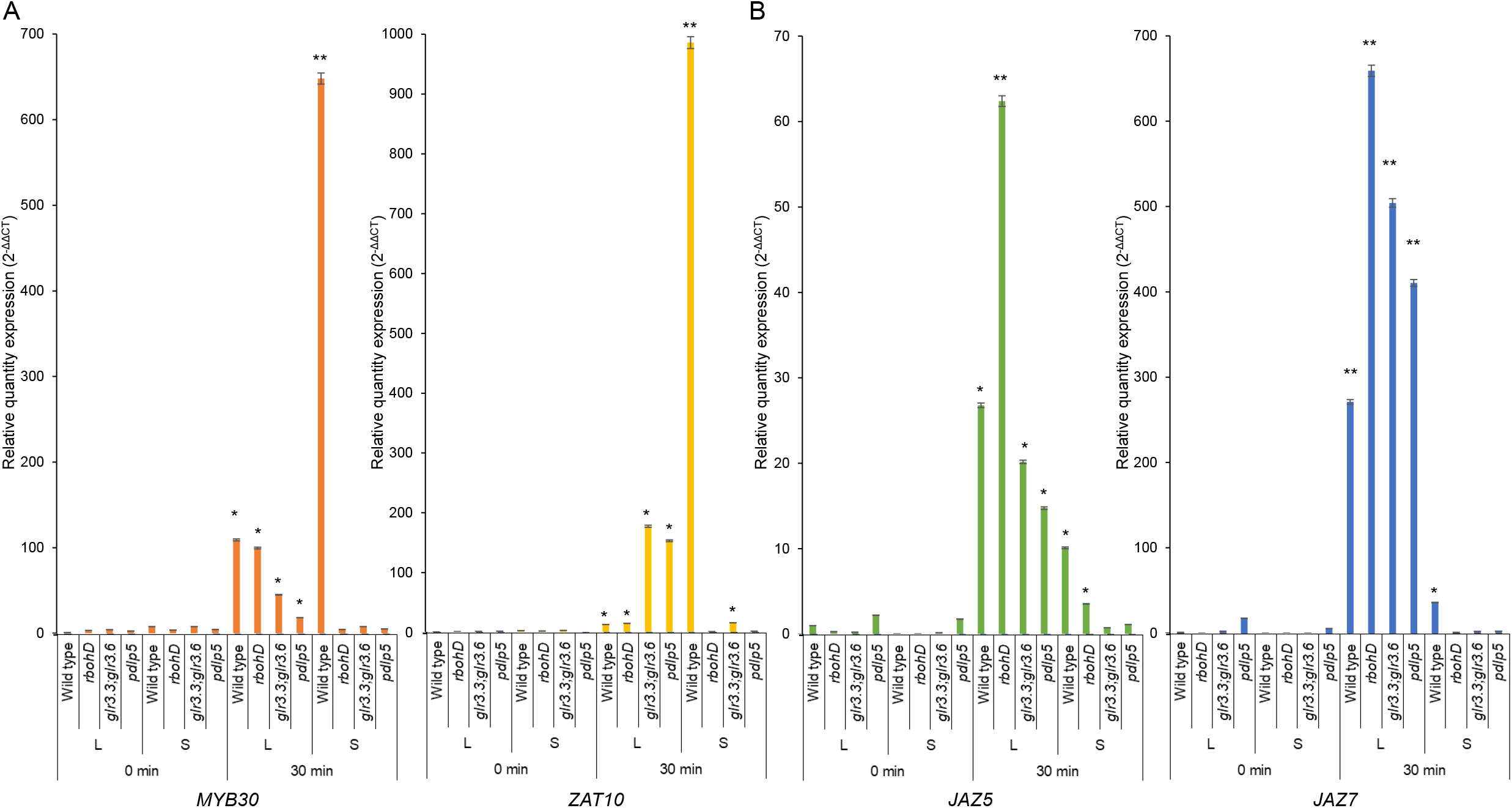
Local and systemic changes in SAA- and SWR-transcript expression in wild type, *rbohD, glr3.3;glr3.6* and *pdlp5* plants subjected to a local wounding or high light stress treatment. (*A*) Real-time quantitative PCR (qPCR) analysis of transcript expression in local and systemic leaves of wild type and *rbohD, glr3.3;glr3.6* and *pdlp5* plants subjected to a local high light (HL) stress treatment applied to a single local leaf. Transcripts tested (*Zat10* and *MYB30*) were previously found to respond to HL stress at the local and systemic leaves of wild-type plants (29, 44, 55). (*B*) Same as (*A*) but plants were subjected to a local wounding treatment applied to of a single local leaf. Transcripts tested (*JAZ5* and *JAZ7*) were shown to respond to wounding at local and systemic leaves of wild-type plants (21, 53). Expression is normalized by internal control (elongation factor 1 alpha) and expression level at 0 min of wild type local leaf. Expression is shown as 2^-ΔΔCT^. Data represents 15 biological repeats for each treatment in each timepoint and 3 technical repeats for each reaction. Student t-test compared to 0 min local leaf of the genotype, SE, N=3, *p < 0.05, **p < 0.01. *Abbreviations used:* glr, glutamate receptor-like; HL, high light; JAZ, jasmonate ZIM-domain protein; PCR, polymerase chain reaction; pdlp5, plasmodesmata localized protein 5; rbohD, respiratory burst oxidase homolog D.

## Discussion

Systemic signaling pathways play a key role in the successful acclimation of plants to rapid changes in their environment (27–29, 53, 55). At least four different signals are thought to mediate systemic signaling in plants in response to abiotic stress and mechanical injury (membrane potential, calcium, ROS, and hydraulic; Figs. 1-4; 4, 6–11). How these signals are interlinked and whether they require each other to propagate, remain however open questions (6–8, 10, 11). Here we reveal that in response to wounding, systemic changes in membrane potential, calcium levels, ROS and hydraulic pressure are coordinated by GLR3.3;GLR3.6 and RBOHD, while in response to HL the RBOHD-mediated systemic ROS signal could be separated from the GLR3.3;GLR3.6-mediated changes in membrane potential and calcium levels (Figs. 1-3, 6). This finding suggests that different stresses could trigger different types of systemic signals that might or might not be co-regulated. This possibility is further supported by our recent findings that during wounding, but not HL stress, the systemic ROS signal can propagate through mesophyll cells (28, 42). Different types of stress could therefore trigger different types of systemic signals that could propagate through different cell layers. This could contribute to the specificity of systemic signaling and convey information regarding the type of stress that triggered the systemic signaling response. Indeed, transcriptomics and SAA studies revealed that the systemic response of plants to wounding, HL or HS is very different, yet all three stresses are thought to trigger rapid systemic membrane potential, calcium, ROS, and hydraulic signals (Figs. 1-4; 6–11, 15, 25–27, 42, 53). It is possible therefore that in response to each specific stress, a specific set of systemic signals is triggered (Fig. 6), and that this set of signals is mediated through the same or different groups of cells. Although the systemic ROS signal may not be linked to systemic calcium and membrane potential signals during HL stress in the *glr3.3;glr3.6* double mutant (Fig. 1; 44), it nevertheless appears to require other signals (perhaps even the systemic calcium and membrane potential signals) to induce a maximal systemic transcript expression response (Fig. 5; 44). The systemic ROS signal, although still propagating in in response to HL stress in the *glr3.3;glr3.6* double mutant was therefore not sufficient to induce a strong systemic transcript expression response (Fig. 5; 44). Further studies are needed to dissect the interactions between the different systemic signals during different stresses, determine how they convey specificity, and how they are linked to different plant hormones such as JA, auxin and ABA.

**Fig. 6.**
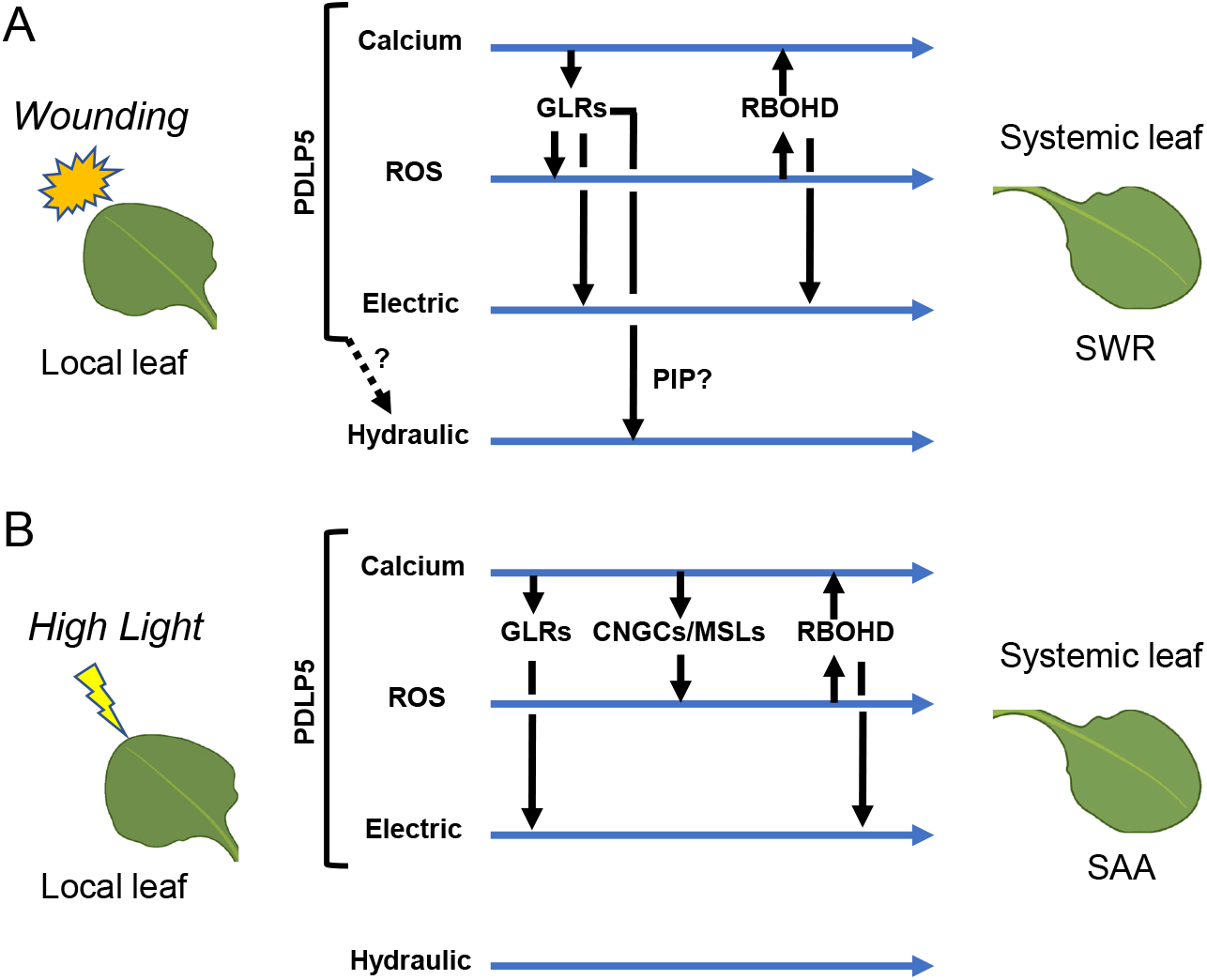
Hypothetical model for the integration of electric, calcium, ROS and hydraulic signals during rapid systemic signaling in plants. (*A*) Wounding of a local leaf is shown to trigger systemic calcium, ROS, electric (membrane potential) and hydraulic signals that trigger a systemic wound response (SWR) in systemic leaves. All systemic signals are shown to be regulated by glutamate receptor-like (GLR) proteins. In addition to GLRs, the respiratory burst oxidize homolog D (RBOHD) protein is shown to regulate ROS and electric signals. The regulation of systemic hydraulic signals is proposed to be mediated by aquaporins (plasma membrane-intrinsic protein; PIP). (*B*) High light (HL) stress applied to a local leaf is shown to trigger systemic calcium, ROS, electric and hydraulic signals that trigger systemic acquired acclimation (SAA) responses in systemic leaves. During systemic responses to HL stress the systemic ROS signal that is dependent on RBOHD can be separated from the systemic calcium, electric and hydraulic signals. GLRs are required for systemic calcium and electric responses but are not linked to systemic hydraulic signals. Instead of GLRs, cyclic nucleotide-gated calcium (CGNCs) and mechanosensitive small conductance–like (MSLs) channels are thought to link the systemic calcium signal with the systemic ROS signal (44). In response to both stimuli (*A* or *B*) plasmodesmata-localized protein 5 (PDLP5) is shown to be required for the systemic calcium, electric and ROS signals. *Abbreviations used:* CGNC, cyclic nucleotide-gated calcium; GLR, glutamate receptor-like; MSL, mechanosensitive small conductance–like; PIP, plasma membrane-intrinsic protein; RBOHD, respiratory burst oxidize homolog D; SAA, systemic acquired acclimation; SWR, systemic wound response.

The findings that the systemic ROS signal can still be detected in the absence of systemic membrane potential and calcium signals during HL stress in the *glr3.3;glr3.6* double mutant (Figs. 1-3), raises the possibility that the systemic ROS signal is not always linked to these signals (Fig. 6). Because the systemic ROS signal appears to be dependent on GLR3.3;GLR3.6 during wounding (Fig. 1), but on CNGC2 and certain MSLs during HL stress (44), it is possible that systemic calcium and membrane potential signals can still occur in the *glr3.3;glr3.6* double mutant during HL stress (possibly mediated by CNGC2 and MSLs), but that they are different in their nature, or much lower in their intensity (compared to those occurring in wild type plants), and that our method may simply cannot detect them. Alternatively, during HL stress the systemic ROS signal may be mediated by a different mechanism that does not require systemic calcium or membrane potential signals, for example through a direct phosphorylation relay between ROS-activated cysteine-rich receptor like kinases and RBOHD, or via direct regulation of RBOHD by redox or nitric oxide (6, 40, 56). An additional difference between HL and wounding is evident by how the hydraulic wave is regulated. During wounding the hydraulic wave requires the function of GLR3.3;GLR3.6, while during HL it is not (Figs. 4, 6). In addition, as opposed to the systemic membrane potential and calcium signals, during HL stress or wounding, the systemic hydraulic signal does not appear to be directly linked to RBOHD (Figs. 4, 6). One intriguing possibility is that systemic hydraulic signals are linked to RBOHF that was recently shown to support the systemic ROS signal, propagating through the vascular bundles of Arabidopsis during systemic responses to HL stress (28). Further studies are of course needed to address this possibility.

One of the most intriguing findings reveled by our study is that in the *glr3.3;glr3.6* double mutant the wound-induced systemic hydraulic signal is suppressed (Fig. 4). Hydraulic signals can be triggered by mechanical injury, rapid changes in stomatal aperture, sudden heat or cold stresses and/or other physical stimuli that will impact the water pressure within the plant vascular bundles (31–33). Why in the *glr3.3;glr3.6* double mutants they are suppressed? Can hydraulic signals be actively regulated by the plant in response to different stimuli? Somewhat akin to how the vascular system of mammalians can contract and affect blood pressure. It is unknown at present whether the phenotype we are observing in the *glr3.3;glr3.6* double mutant (Fig. 4B) is an outcome of changes in stomata and/or PD reactions, changes in ion fluxes or pH, and/or other unknown at present mechanisms that shape or block the systemic hydraulic signal and require GLR3.3;GLR3.6. One possible mechanism could be mediated through the function of aquaporins that are actively regulated in plants by phosphorylation, calcium, and/or other reactions (33, 57–60). If for example a particular signal such as wounding would trigger a GLR3.3;GLR3.6-dependent signaling mechanism that will cause the simultaneous closure of aquaporins in cells, at and around the vascular bundle, this process could cause an increase in hydraulic pressure in the xylem vessels; because the pressure that is usually relieved from the xylem column into neighboring cells will now cease to be relieved. A similar effect would occur for example in a long water pipe that feeds many open taps around a city block. If all taps are suddenly and simultaneously closed, the water pressure will build up. Because hydraulic signals were proposed to play an important role in systemic signaling, triggering systemic calcium signals through different mechanosensory proteins (4, 6, 8), the suppression of the systemic hydraulic signal in the *glr3.3;glr3.6* double mutant could play an important signaling role. Further studies are of course needed to address these intriguing possibilities and to determine how GLR3.3;GLR3.6 regulate systemic hydraulic signals.

In addition to aquaporins, PD can also play an important role in mediating different systemic signals. It was recently found that PD functions, mediated through the PDPL1 and 5 proteins, are important for the propagation of the systemic ROS signal during systemic responses to HL stress (44). Here we show that PDLP5-regulated functions are also essential for systemic membrane potential and calcium responses (Figs. 2, 3). In addition to the apoplast, that plays a major role in mediating systemic signals between cells (6–9, 11, 61), PD may therefore also play a canonical role in mediating rapid systemic responses in plants (Figs. 1-3; 44). Because PD connect the plasma membrane of one cell to another and mediate the transfer of small molecules between cells (62), their function could be critical for the rapid cell-to-cell spread of systemic membrane potential and calcium signals. During HL stress PDLP1 and 5 were recently shown to regulate the rapid opening of PD pores and to be required for the systemic signal to propagate through the plant and activate SAA in systemic tissues (44). A similar function during wounding could therefore facilitate the systemic calcium, ROS, and membrane potential signals in response to wounding. Further studies are of course required to determine the role of PD in the rapid systemic signaling process of plants.

The whole-plant live ROS imaging method we developed (26) has been validated in different studies using different mutants (27, 44, 55), omics tools (27, 29, 41), and more recently whole-plant imaging of transgenic plants with stable expression of cytosolic reduction-oxidation sensitive green fluorescent protein 1 (roGFP1; 40). Although the whole-plant live imaging methods, presented in our current study to record systemic changes in calcium levels and membrane potentials (Figs. 2, 3), would need to be followed by more detailed studies of electric wave measurements using different electrodes (to determine the exact types of electric waves involved; 7, 19), as well as using transgenic plants with stable expression of different ratiometric sensor proteins for calcium measurements in different subcellular compartments (to better characterize the calcium waves involved; 21), our findings that systemic changes in calcium and membrane potential are abolished in the *glr3.3;glr3.6* double mutant that was shown to display suppressed electric and calcium waves in response to wounding (16–18, 21); similar to how the ROS wave is suppressed in the *rbohD* mutant in response to HL stress (Fig. 1; 25, 26, 28), demonstrate that the new methods presented here can be used to study the impact of specific mutations on different rapid whole-plant systemic signaling processes. Of course, as indicated above, further studies are needed to dissect and determine the exact identity of the different systemic signals involved in these processes. Nonetheless, the results presented in Figs. 1-3 demonstrate that all 3 methods for live imaging of systemic changes in ROS, calcium and membrane potential adhere to what is previously known to occur in the *glr3.3;glr3.6* and *rbohD* mutants during wounding or HL stress, respectively (16–18, 21, 25–28).

Taken together, our findings reveal that different systemic signaling mechanisms and pathways could be activated by different stresses, that many of these systemic signaling pathways, including hydraulic pressure waves are interlinked via the function of GLRs, and that many of them require the function of PD-associated proteins (Fig. 6). The web of rapid systemic signals propagating through the plant vascular system during different stresses could therefore transmit different types of signals in response to different stresses, and these may or may not be linked with each other, depending on the type of stress triggering them.

## Materials and Methods

### Plant material, growth, and stress treatments

Homozygous *Arabidopsis thaliana* wild type (Col-0) and knockout *rbohD* (AT5G47910; 63), *glr3.3;glr3.6* (AT1G42540, AT3G51480; 16, 17), and *pdlp5* (AT1G70690; 44) plants were germinated and grown on peat pellets (Jiffy-7, Jiffy International, Kristiansand, Norway) under controlled conditions of 10hr/14hr light/dark regime, 50 μmol photons s^-1^ m^-2^ and 21°C for 4 weeks (44). Plants were subjected to HL stress by illuminating a single leaf with 1700 μmol photons s^-1^ m^-2^ using a ColdVision fiber optic LED light source (Schott, Southbridge, MA, USA), or to wounding by puncturing a single leaf, with 18 dressmaker pins (Singer, Murfreesboro, TN, USA), as described earlier (26–28, 40, 44, 55).

### Whole-plant fluorescence imaging of ROS, calcium, and membrane potential

For ROS imaging, plants were fumigated with 50 μM H_2_DCFDA (Millipore-Sigma, St. Louis, MO, USA) for 30 min in a glass container using a nebulizer (Punasi Direct, Hong Kong, China), as described in (26). Similarly, for calcium imaging, plants were fumigated with 4.5 μM Fluo-4-AM (Becton, Dickinson and Company, Franklin Lakes, NJ, USA; 45–47), and for membrane potential imaging, plants were fumigated with 20 μM DiBAC4(3) (Biotium, Fermont, CA; 49–52, 64), in 1.5 mM KCl buffer for 30 min. Following fumigation, local HL stress or wounding were applied to a single leaf as described above. Fluorescence images (ex./em. 480 nm/ 520 nm) were then acquired using IVIS Lumina S5 (PerkinElmer, Waltham, MA, USA) for 30 min as described in (26). Fluorescence accumulation was analyzed using Living Image 4.7.2 software (PerkinElmer) using the math tools (26). Time course images were generated and radiant efficiency of regions of interest (ROI) were calculated (26). Each data set includes standard error of 8-12 technical repeats and a Student t-test score (26). Dye penetration controls were performed by fumigation of plants with 1 mM hydrogen peroxide for 10 min following the H_2_DCFDA fumigation (for ROS; *SI Appendix*, Fig. S1; 26); fumigation of plants with 1 mM CaCl_2_, 10 μM calcium ionophore A23817 (Millipore-Sigma, St. Louis, MO, USA; 65) for 10 min following the Fluo-4 AM fumigation (for intracellular calcium imaging; *SI Appendix*, Fig. S2), or fumigation of plant with 20 μM sodium cholate (49) for 10 min following the DiBAC4(3) fumigation (for membrane potential; *SI Appendix*, Fig. S3).

### Confocal Microscopy

Plants were fumigated with H_2_DCFDA, Fluo-4-AM or DiBAC4(3) for 30 min and detached leaves were imaged with a confocal microscope as described in (26). H_2_DCFDA, Fluo-4-AM and DiBAC4(3) z-stacks were generated using Leica TCS SP8 MP (×20 magnification), 8% laser intensity (excitation/emission 495 nm/520 nm); z-stacks-composed 3D projections from 35-45 slices of 0.4 μm were generated with Leica Application Suit X. Images were acquired and analyzed at the University of Missouri Molecular Cytology Core facility.

### Hydraulic pressure measurements

Changes in systemic leaf turgor pressure following wounding or HL stress applied to a single local leaf were recorded using the ZIM-probe system (Yara International ASA, Oslo, Norway; 54). In short, a single systemic leaf was connected to 2 magnetic probes that included a pressure sensor between them (54). The turgor pressure force against the magnetic pressure was recorded and transmitted to a receiver every 60 sec. Following magnetic probe attachment, the system was allowed to stabilize for 3 hours and a HL stress or wounding treatment was applied to a single local leaf. Measurements were then conducted for an additional 30 min following the stress treatment. Untreated plants were similarly measured as controls, but without application of HL stress or wounding (*SI Appendix*, Fig. S4). Results are calculated as percent of control, which is the measured pressure in the leaf 30 min prior to the stress application. All experiments were performed 4 hours following the start of the day photoperiod. Each data set includes standard error of 5-10 biological repeats.

### Local and systemic SAA and SWR transcript expression

To measure the transcriptional response of local and systemic leaves to HL or wounding stress in 4-week-old plants, HL or wounding were applied to a single leaf for 30 min. Exposed leaf (local) and unexposed fully developed younger leaf (systemic) were collected for RNA extraction. RNA was extracted using Plant RNeasy kit (Qiagen, Hilden, Germany) according to the manufacture instructions. Quantified total RNA was used for cDNA synthesis (PrimeScript RT Reagent Kit; Takara Bio, Kusatsu, Japan). Transcript expression was quantified by real-time qPCR using iQ SYBR Green supermix (Bio-Rad Laboratories, Hercules, CA, USA), as described in (44), with specific primers for HL: *ZAT10* (AT1G27730) 5′- ACTAGCCACG TTAGCAGTAGC-3′ and 5′- GTTGAAGTTTGACCGGAAGTC-3′ and *MYB30* (AT3G28910) 5′- CCACTTGGCGAAAAAGGCTC-3′ and 5′- ACCCGCTAGCTGAGGAAGTA-3′. For wounding (21), *JAZ5* (AT1G17380) 5′- TCATCGTTATCCTCCCAAGC-3′ and 5′- CACCGTCTGATTTGATATGGG-3′ and *JAZ7* (AT2G34600) 5′- GATCCTCCAACAATCCCAAA-3′ and 5′- TGGTAAGGGGAAGTTGCTTG-3′. *Elongation factor 1 alpha* (5′- GAGCCCAAGTTTTTGAAGA-3′ and 5′-TAAACTGTTCTTCCAAGCTCCA-3′) was used for normalization of relative transcript levels. Primer efficiency was at the 0.99-1.04 range (44, 55). Results in the exponent of base 2 delta-delta terminal cycle were obtained by normalizing the relative transcript and comparing it to control wild type from local leaf. The data represents 15 biological repeats and 3 technical repeats for each reaction. Standard error and Student t-test were calculated with Microsoft Excel.

### Statistical analysis

Statistical analysis for ROS, calcium and membrane potential changes (total radiant efficiency), and real-time quantitative PCR transcript expression was performed by two-sided student t-test and results are presented as mean ± SE, *p < 0.05, **p < 0.01, ***p < 0.001. Hydraulic pressure results are presented as mean ± SE.

## Author Contributions

Y.F. performed experiments and analyzed the data. R.M. and Y.F. designed experiments, analyzed the data and wrote the manuscript.

## Acknowledgments

We thank Professor E. Farmer, University of Lausanne for seeds of the *glr3.3;glr3.6* double mutant. This work was supported by funding from the National Science Foundation (IOS-1353886, MCB-1936590, IOS-1932639) and the University of Missouri.

## *SI Appendix* Figure Legends

**Fig. S1.**
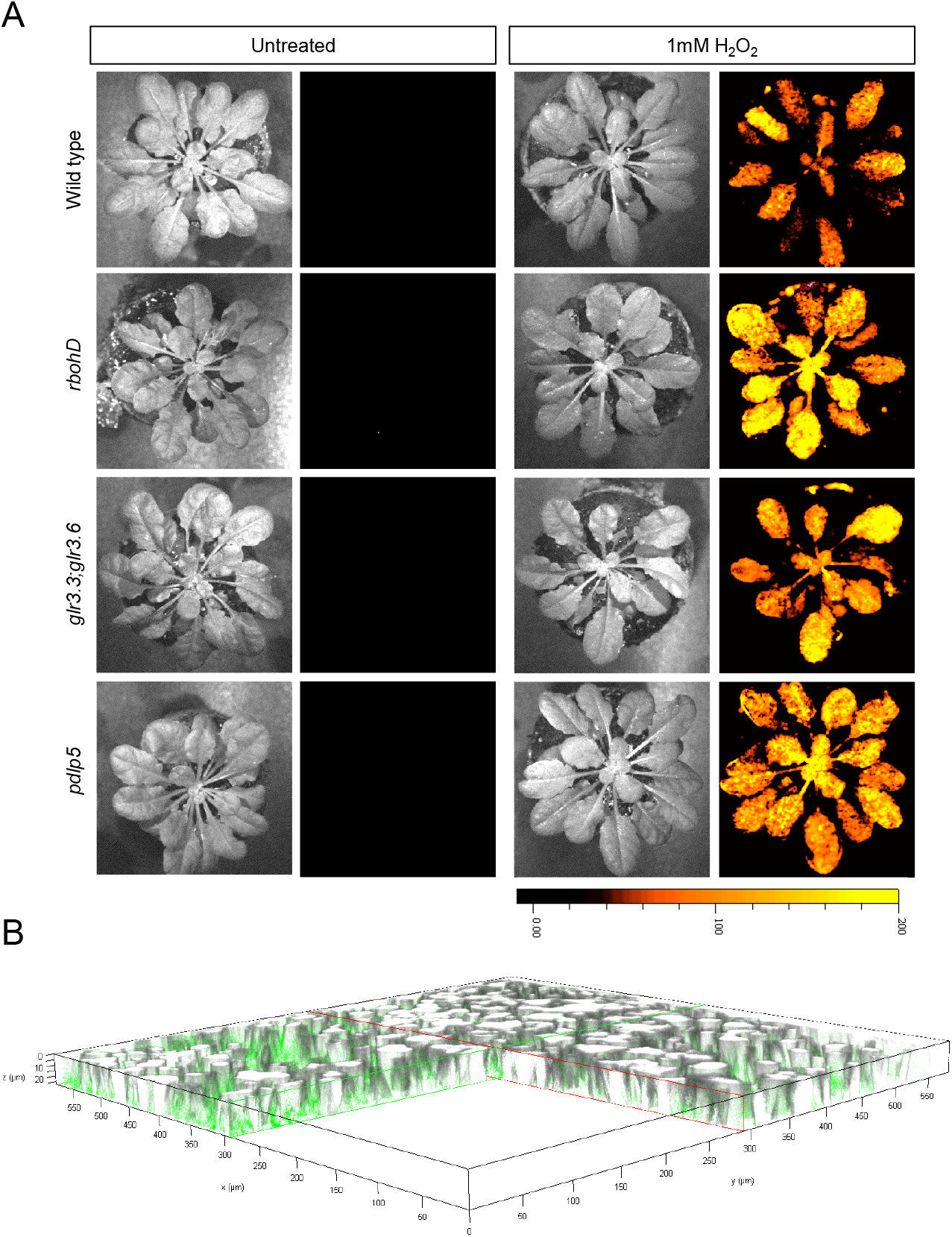
Whole-plant ROS imaging following H_2_O_2_ fumigation of wild type plants and the different mutants. (*A*) To control for dye penetration and function in the different mutants, plants were fumigated with H_2_DCFDA for 30 min and then with 1 mM H_2_O_2_ (to mimic enhanced ROS accumulation), for 10 min, as described in (26). Images shown are representative of 3 independent experiments. (*B*) To determine dye penetration into leaves, plants were fumigated with H_2_DCFDA for 30 min and detached leaves were imaged with a confocal microscope. Representative three-dimensional projection of Z-stacked confocal images of H_2_DCFDA-fumigated wild type leaves (fluorescent overlay 3D) are shown. Images shown are representative of 3 independent experiments.

**Fig. S2.**
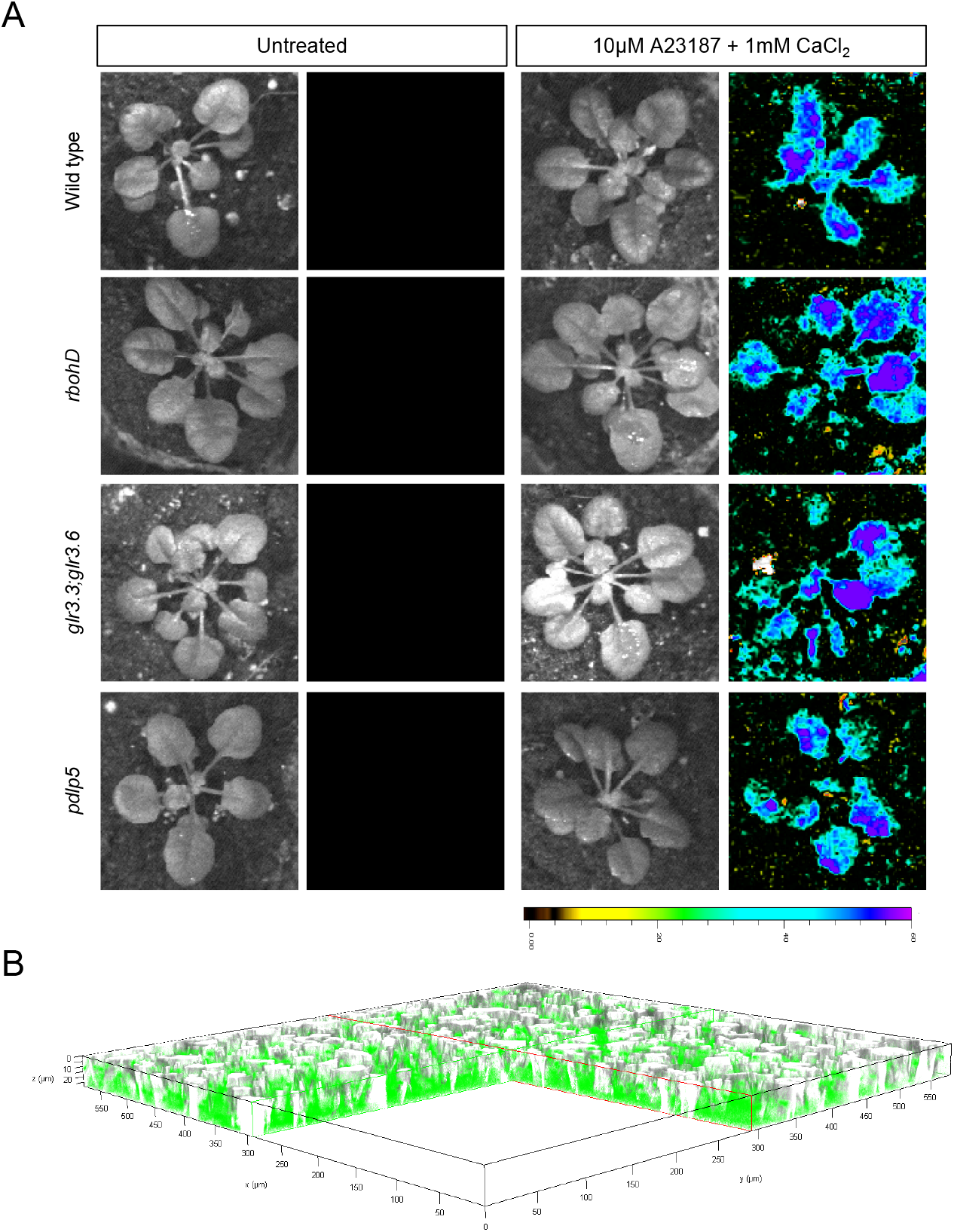
Whole-plant calcium imaging following fumigation of wild type plants and the different mutants with 1 mM CaCl_2_ and 10 μM calcium ionophore A23817. To control for dye penetration and function in the different mutants, plants were fumigated with Fluo-4 AM for 30 min and then with 1 mM CaCl_2_, 10 μM calcium ionophore A23817 (to mimic enhanced intracellular calcium levels), for 10 min. Images shown are representative of 3 independent experiments. (*B*) To determine dye penetration into leaves, plants were fumigated with Fluo-4 AM for 30 min and detached leaves were imaged with a confocal microscope. Representative three-dimensional projection of Z-stacked confocal images of Fluo-4 AM-fumigated wild type leaves (fluorescent overlay 3D) are shown. Images shown are representative of 3 independent experiments.

**Fig. S3.**
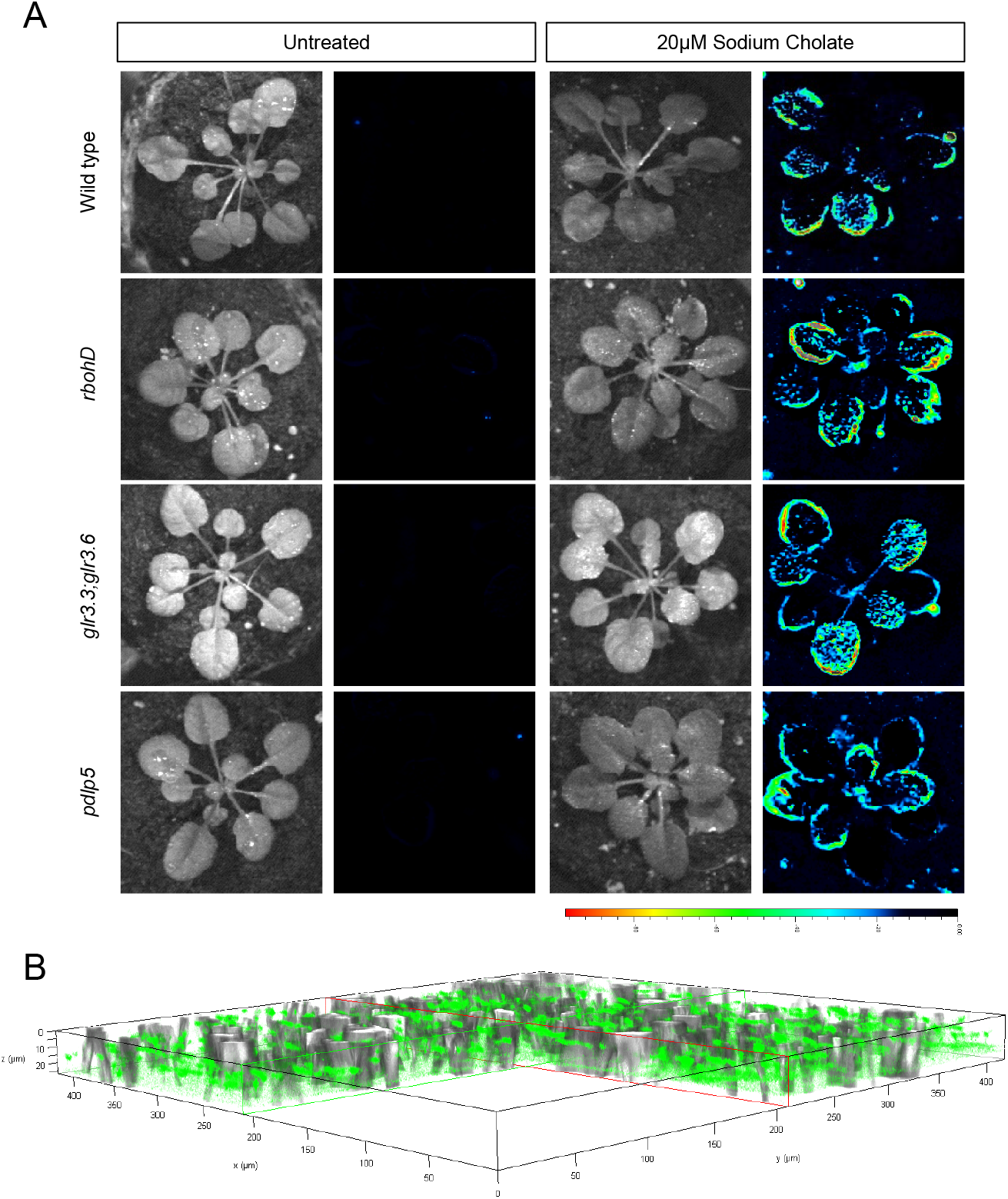
Whole-plant membrane potential imaging following fumigation of wild type plants and the different mutants with 20 μM sodium cholate. To control for dye penetration and function in the different mutants, plants were fumigated with DiBAC4(3) for 30 min and then with 20 μM sodium cholate (to mimic changes in membrane potential), for 10 min. Images shown are representative of 3 independent experiments. (*B*) To determine dye penetration into leaves, plants were fumigated with DiBAC4(3) for 30 min and detached leaves were imaged with a confocal microscope. Representative three-dimensional projection of Z-stacked confocal images of DiBAC4(3)-fumigated wild type leaves (fluorescent overlay 3D) are shown. Images shown are representative of 3 independent experiments.

**Fig. S4.**
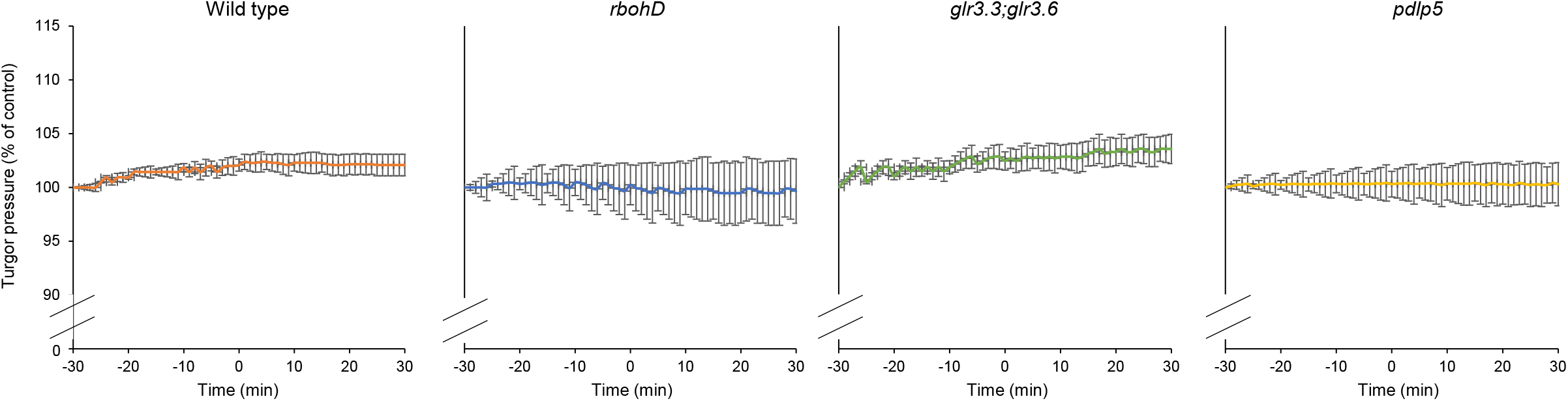
Systemic changes in hydraulic pressure in untreated wild type, *rbohD, glr3.3;glr3.6* and *pdlp5* plants (control for Fig. 4). Continues systemic leaf turgor pressure measurements of wild type, *rbohD, glr3.3;glr3.6* and *pdlp5* plants, from 30 min prior (−30 min) to 30 min post (30 min) untreated (equivalent timing to treatments of high light stress or wounding of a single local leaf; Fig. 4) are shown. Hydraulic pressure is represented as percent of the initial measured turgor pressure at −30 min. All experiments were repeated at least 5 times. SE. *Abbreviations used:* glr, glutamate receptor-like; HL, high light; pdlp5, plasmodesmata localized protein 5; rbohD, respiratory burst oxidase homolog D.

